# Comparative multivariate decoding adjudicates theories of semantic representation in the anterior temporal lobes and the rest of the cortex

**DOI:** 10.1101/2025.08.22.671718

**Authors:** Saskia L. Frisby, Christopher R. Cox, Ajay D. Halai, Matthew A. Lambon Ralph, Timothy T. Rogers

## Abstract

The anterior temporal lobes (ATLs) are known to support semantic cognition, but theories about their precise representational coding vary. We collected 7T-fMRI data with a novel acquisition sequence designed to improve signal quality in the ATLs, then employed a pioneering analytical approach (comparative multivariate decoding) to adjudicate between theories. Specifically, we applied multiple decoding methods, each making different assumptions about the content, nature or location of representations within the ATLs, then used the pattern of results *across* methods to adjudicate competing hypotheses. The results suggest that the ATLs represent domain-general semantic information via a multidimensional inchoate vector-space code that is anatomically clustered within and across individuals, and that posterior temporal and occipitotemporal regions utilise a similar domain-general, inchoate vector-space code. More generally, the comparative multivariate analytical framework utilised here has the potential to reveal how the brain represents, not just semantic knowledge, but any kind of information.

## 1. Introduction

Humans have a remarkable ability to understand the world around them – to recognise objects and make inferences about their properties, to comprehend and produce language, and to predict and influence events. These capacities, collectively known as *semantic cognition*, rely on neural systems that remain poorly understood, with controversy surrounding two general questions. The first concerns the role of the anterior temporal lobes (ATLs) – regions known to support semantic cognition ^1–10^ but which are the subject of differing views about their function and the representations they subserve. The second concerns regions beyond the ATLs – which ones are also important for semantic cognition, what information they encode, and whether their functions differ from the ATL. These divergent views may arise because functional neuroimaging studies of semantics apply different analytic techniques, each adopting different assumptions about the neural code that limit the kinds of representation it can and cannot detect ^11^. Because most studies adopt a single kind of analysis, each is unable to adjudicate between theories of how neural activity encodes semantic information.

To address this challenge, we introduce *comparative multivariate decoding* (cMVD) as an approach for testing competing theories of representation in the brain. From the literature we first delineate several distinct hypotheses about how neural activity may encode information of interest. We then choose a set of multivariate decoding models such that each hypothesis about the neural code predicts a unique pattern of outcomes *across* decoding models. After using each model type to decode information from a given dataset, we compare the pattern of successes and failures to those predicted by the different hypotheses, thereby adjudicating them.

While model comparison is a backbone of statistics generally and some prior imaging studies have compared the accuracy of different multivariate methods^12,13^, cMVD differs from earlier work by drawing explicit connection between the representational hypotheses to be adjudicated and the family of decoding models selected for comparison. Rather than just comparing the respective accuracies of various decoding approaches, we instead consider what kinds of neural signal each approach can/cannot detect and, critically, how these signal-detection properties relate to the competing representational hypotheses. The results subsequently establish *what*, *how*, and *where* information of interest is encoded in functional brain activations, in a manner that goes beyond single-method approaches. The current paper uses cMVD to resolve open questions about semantic representation.

### 1.1. Delineating representational hypotheses

Whilst many sources of evidence converge to implicate the ATL in semantic representation (e.g. clinical neuropsychology ^3,7,14^, intracranial electrophysiology ^6,15,16^, functional neuroimaging ^5,17^, neurostimulation ^18^, computational modelling ^2,4^ ^19^), hypotheses about its precise role differ along three different dimensions:

#### 1. Domain of representation (“what”)

From evidence of category-specific patterns of semantic impairment and functional activation ^7,10^, some hypotheses propose that semantic representations in the ATL are *animal-specific* ^8,9^ whereas other regions support knowledge about manmade objects ^20^, social and emotional concepts ^21^, or other categories ^22^. Alternatively, seemingly category-specific patterns may largely reflect confounding factors ^7,23,24^ that, once controlled, reveal the ATLs to support semantic knowledge for all conceptual domains ^25,26^.

#### 2. Nature of representation (“how”)

The literature offers three different proposals about the nature of semantic representations within the ATLs: *2A. Pointers.* The ATLs may contain “pointers” (within distinct “convergence zones”) for different concepts, each binding together its associated features encoded elsewhere in the cortex ^27^. For instance, the *robin* pointer may connect properties such as *has feathers, has a red breast, can fly* etc. Under this hypothesis, perception of the word “robin” activates the *robin* pointer in the ATL, which then activates the robin’s shape, pattern of motion, typical colour, other verbal descriptors, etc. in various remote cortical regions. Each concept has its own pointer—for instance, the robin pointer connects properties of robins; the raven pointer connects properties of ravens; and the raven and robin pointers are separate, individuated entities within the convergence zone, even though robins and ravens share many properties encoded elsewhere in cortex. Consequently, on this view it should not be possible to decode graded semantic similarity relations between items from the ATL pointers alone.

#### 2B. Semantic features

Different neural populations within the ATL (and elsewhere in cortex) may each *independently* encode an interpretable semantic feature dimension, such as animacy (animate/inanimate), size (large/small), movement (can fly/can’t fly), and so on ^28,29^. Semantic features capture graded conceptual similarity relations ^30,31^ (i.e. more similar concepts have more features in common) so, on this view, ATL activity patterns should express such similarities within the domain represented. Because different semantic dimensions are encoded by distinct neural populations, however, fitting a model to decode one dimension (e.g. animacy) should not aid in finding signal important for another orthogonal dimension (e.g. type of movement). Thus decoding performance should be best when *different* models are fitted to decode each dimension.

#### 2C. Inchoate semantic vector spaces

Neural populations within the ATL may *conjointly* encode a graded semantic representation space, without each population corresponding to an interpretable dimension. On this view, ATL activity patterns express graded conceptual similarities between stimuli within the domain of representation, but each semantic dimension (animacy, size, type of movement, etc) is encoded by patterns over multiple populations, and each population partially encodes information about multiple different semantic dimensions ^1,2,4,32–38^. Thus fitting a model to decode one semantic distinction (e.g. animacy) *can* aid in finding signal relevant to other orthogonal distinctions (e.g. type of movement), and decoding models should perform better when constrained to decode multiple semantic dimensions from the *same* neural populations. Following Rogers & McClelland^38^, we refer to this hypothesis as an *inchoate* semantic vector space.

### 3. Location of the representation (“where”)

Functional brain imaging often assumes that the neural populations representing semantic information are contiguous within individual brains and that the semantic system is organised similarly across individuals. Recent applications of multivariate decoding suggest, however, that semantic representations can be anatomically dispersed within and variable across individuals ^29,39^. Thus hypotheses vary in their localization of representations within the ATL. If representations are anatomically clustered (i.e. loosely contiguous within and similarly located across individuals), then decoding models constrained to prefer anatomically clustered representations should outperform models that ignore anatomical contiguity and similarity.

The combination of these three aspects leads to 12 distinct hypotheses about the nature and location of semantic representations in the ATL, listed in Table 1.

### 2. Selecting a set of models for cMVD

With cMVD we aim to adjudicate these hypotheses by designing a family of decoding analyses across which each hypothesis predicts a *different pattern* of relative performance. For instance, pointer-based representations predict that it should not be possible to decode graded similarities between items within or outside the domain of representation, while animal-specific hypotheses predict that the ATLs only support knowledge about animals. If ATL representations are both pointer-based and animal-specific, a binary classifier fitted to ATL activations should succeed at discriminating animate from inanimate stimuli (since only animals activate a pointer), but a regression model predicting between-animal semantic distances should fail (since pointers do not encode graded semantic similarities). In contrast, a domain-general pointer representation should *fail* for both decoding goals (since animals and non-animals both activate ATL pointers); an animal-specific feature-based representation should achieve both goals but fail at decoding between-item similarities for non-animals (since non-animal features are not encoded within ATL); and a domain-general feature-based representation should achieve all three goals (binary classification, within-animate similarities, within-inanimate similarities). Following this logic, the 12 hypotheses can be distinguished by comparing the performance of decoding models that vary three independent factors:

**Table 1.**
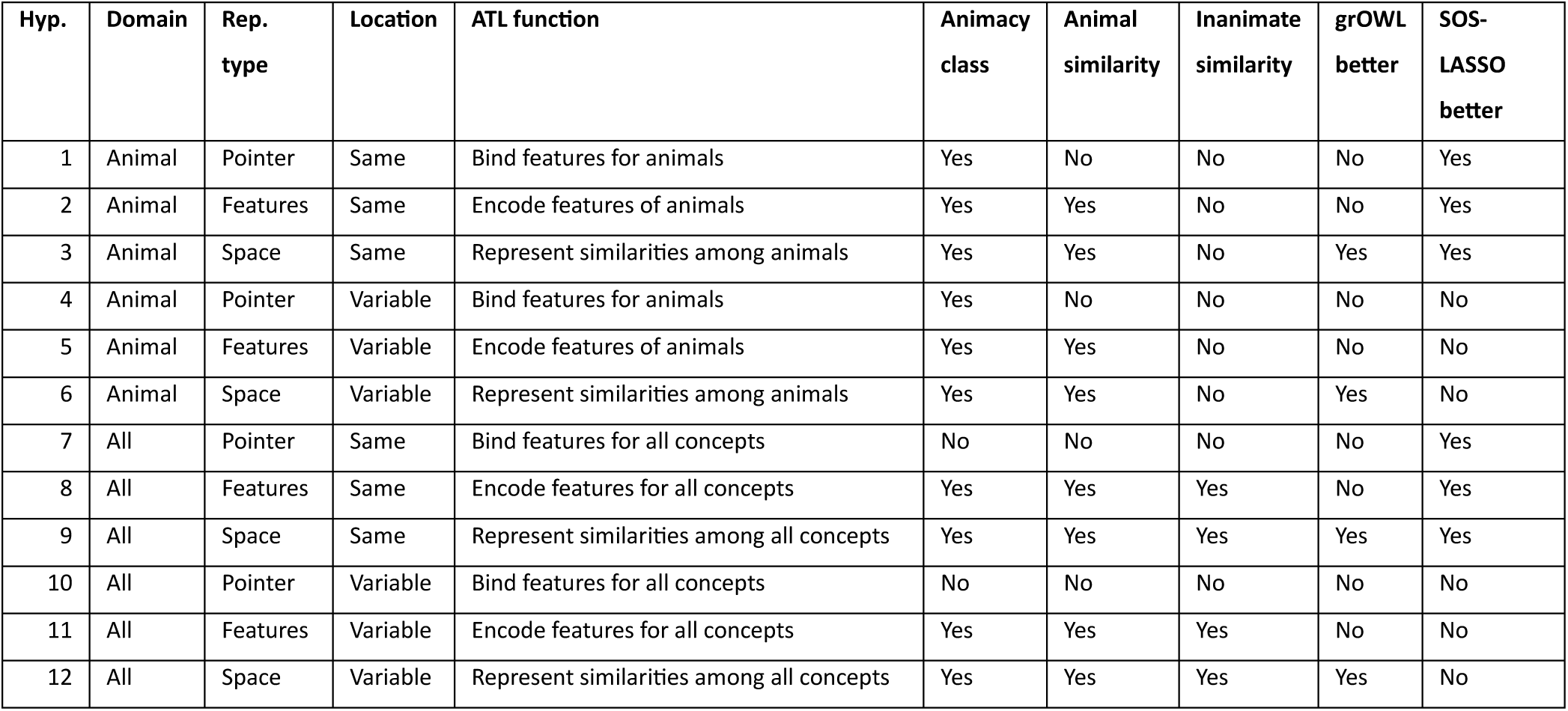
Pattern of results predicted by each of 12 hypotheses across five comparative decoding analyses. Hyp.: Hypothesis, Rep. type: representation type. **Animacy class:** Can classifiers successfully predict stimulus animacy? **Animal similarity:** Can RSL models successfully predict multidimensional semantic similarities between animate items? **Inanimate similarity:** Can RSL models successfully predict multidimensional semantic similarities among inanimate items? **grOWL better:** Do RSL models predicting graded semantic structure perform better with grOWL regularization (which prefers solutions that predict multiple target dimensions from the same neural features) than with LASSO regularization (which does not have this preference)? **SOS-LASSO better:** Do classifiers perform better with SOS-LASSO regularization (which encourages solutions where predictive features are anatomically clustered) than with LASSO regularization (which does not have this preference)?

| Hyp. | Domain | Rep. type | Location | ATL function | Animacy class | Animal similarity | Inanimate similarity | grOWL better | SOS-LASSO better |
| --- | --- | --- | --- | --- | --- | --- | --- | --- | --- |
| 1 | Animal | Pointer | Same | Bind features for animals | Yes | No | No | No | Yes |
| 2 | Animal | Features | Same | Encode features of animals | Yes | Yes | No | No | Yes |
| 3 | Animal | Space | Same | Represent similarities among animals | Yes | Yes | No | Yes | Yes |
| 4 | Animal | Pointer | Variable | Bind features for animals | Yes | No | No | No | No |
| 5 | Animal | Features | Variable | Encode features of animals | Yes | Yes | No | No | No |
| 6 | Animal | Space | Variable | Represent similarities among animals | Yes | Yes | No | Yes | No |
| 7 | All | Pointer | Same | Bind features for all concepts | No | No | No | No | Yes |
| 8 | All | Features | Same | Encode features for all concepts | Yes | Yes | Yes | No | Yes |
| 9 | All | Space | Same | Represent similarities among all concepts | Yes | Yes | Yes | Yes | Yes |
| 10 | All | Pointer | Variable | Bind features for all concepts | No | No | No | No | No |
| 11 | All | Features | Variable | Encode features for all concepts | Yes | Yes | Yes | No | No |
| 12 | All | Space | Variable | Represent similarities among all concepts | Yes | Yes | Yes | Yes | No |

### 1. The decoding target

Models can be fitted to categorise stimuli as animate/inanimate or to predict graded similarities among stimuli on multiple semantic dimensions within and across animate/inanimate domains. For classification we can use regularised logistic regression. For predicting graded similarities we can use representational similarity learning (RSL), an extension of regularised linear regression that fits decoding models to predict target coordinates on multiple dimensions simultaneously ^16,40^.

### 2. Regularization across dimensions

When decoding graded similarities, we can fit models that predict each semantic dimension independently (regularised with a standard loss such as LASSO a.k.a. the L1 norm), or that prefer solutions in which the same neural populations are used to predict all dimensions simultaneously (regularised with group ordered weighted LASSO or grOWL^16,40^).

### 3. Regularization across space

For binary classification, we can fit models that ignore the anatomical location of features, or that prefer solutions that are anatomically clustered within and across participants (sparse-overlapping-sets LASSO or SOS-LASSO^41^).

Table 1 indicates how each of the 12 hypotheses predicts a different pattern of comparative outcomes across these analyses:

The analyses selected have three important properties for cMVD. First, each directly connects to a specific hypothesis about the nature of the neural representation—for instance, logistic regression is designed to find a binary categorical code and thus connects to domain-specific hypotheses; grOWL regularization is designed to find solutions where all target dimensions are encoded across the same features and thus connects to inchoate vector-space hypotheses; SOS-LASSO regularization is designed to prefer solutions where features reside in similar locations within and across individuals and thus connects to hypotheses about consistently localised representation; etc. The relative success of each approach thus provides clues about the true nature of the neural signal. Second, all approaches fall within the same statistical framework of regularised regression, requiring similar procedures for model fitting and assessment and yielding solutions in the form of model coefficients that weight voxel activations to predict target structure. Thus the various solutions are directly comparable to one another in their performance metrics (i.e., decoding accuracy on held-out items) and in the solutions they yield (i.e., comparison and analysis of distributions of selected voxels, that is, those receiving non-zero weights in the decoder). Third, the set of analyses together are minimally sufficient to discriminate the specified hypotheses as shown in Table 1. Thus while the literature contains many other potential decoding approaches (e.g. encoder/decoder^29,42^; representational similarity analyses^43,44^; comprehensively reviewed elsewhere^11^), our cMVD approach excludes them because each requires somewhat different fitting and evaluation methods and they do not aid in further adjudicating the denoted hypotheses.

Outside the ATL, the same comparative approach can be used to decode whole-cortex neural patterns, yielding information about where and how other cortical systems encode semantic information. For instance, by comparing classification to RSL we can assess whether any brain areas encode binary animacy information without also encoding semantic similarity structure; by comparing RSL regularised with LASSO vs. grOWL, we can assess whether distinct semantic dimensions are encoded by different or overlapping neural populations outside the ATL; and by contrasting results of individual vs. SOS-LASSO we can assess whether semantic information is anatomically clustered. Note that nothing precludes different hypotheses (Table 1) being true in different regions.

We applied this comparative approach to new data collected using 7T-fMRI, which produces a greater signal-to-noise ratio (tSNR) than 3T-fMRI and so is advantageous for decoding. We mitigated the magnetic susceptibility effects that affect ATL imaging by using a new multi-echo multi-band scanning protocol recently shown to improve functional contrast-to-noise and to boost decoding accuracy from the ATL at 7T whilst maintaining signal quality across the rest of the brain[FrisbyETAL2026]. We focused first on adjudicating hypotheses of ATL function, then considered implications for semantic representation in the rest of the cortex. The results illuminate the neural bases of semantic representation and suggest a new approach for resolving conflicting patterns of results in cognitive neuroscience more broadly.

## 2. Results

### Data and approach

7T-fMRI data were collected from 27 participants as they named 100 line drawings of familiar items, half animate and half inanimate. We fit a first-level (within-participant) general linear model (GLM) to each participant’s data and extracted one whole-brain beta image per item, taking the beta value as an estimate of the voxel’s BOLD response to the item. These values constituted the “features” passed to the decoder models.

To adjudicate hypotheses about ATL function, we constructed a ventral temporal lobe (vTL) region of interest (ROI) in the left and right hemispheres based on previous electrocorticography (ECoG) studies^16,45,46^ suggesting that cortical surface voltages across this region encode semantic information. For greater anatomical specificity we further subdivided the ROI in each hemisphere into anterior and posterior sections. Because ATL cytoarchitecture features gradual rather than sharp transitions^47^ with no clear boundary between anterior and posterior we simply bisected the full region along its anterior-posterior axis. We then analysed the anterior and posterior halves independently as well as the whole ROI, separately for each hemisphere (6 total ROIs). For the whole-cortex analysis, we applied a cortical grey matter mask. Data from all voxels within the ROI or mask were provided as input to the decoding models.

We assessed binary classification (animate/inanimate) using logistic regression models fit with LASSO regularization (which prefers sparse solutions, i.e. only a few of the available features encode the target information) or with SOS-LASSO (which prefers solutions that are both sparse and anatomically clustered within and across participants).

We used RSL to predict each item’s coordinates on three orthogonal semantic dimensions derived from a set of semantic feature norms ^16,48^ (see Figure 1). The first component (which accounts for 81.1% of the variance) separates animate from inanimate items and also somewhat differentiates animal subclasses. Significant decoding of this dimension is compatible both with a neural code representing independent features that systematically covary with animacy (e.g., *can move*, *has eyes*, etc.) or a neural code that represents a direction within a semantic vector space.

**Figure 1:**
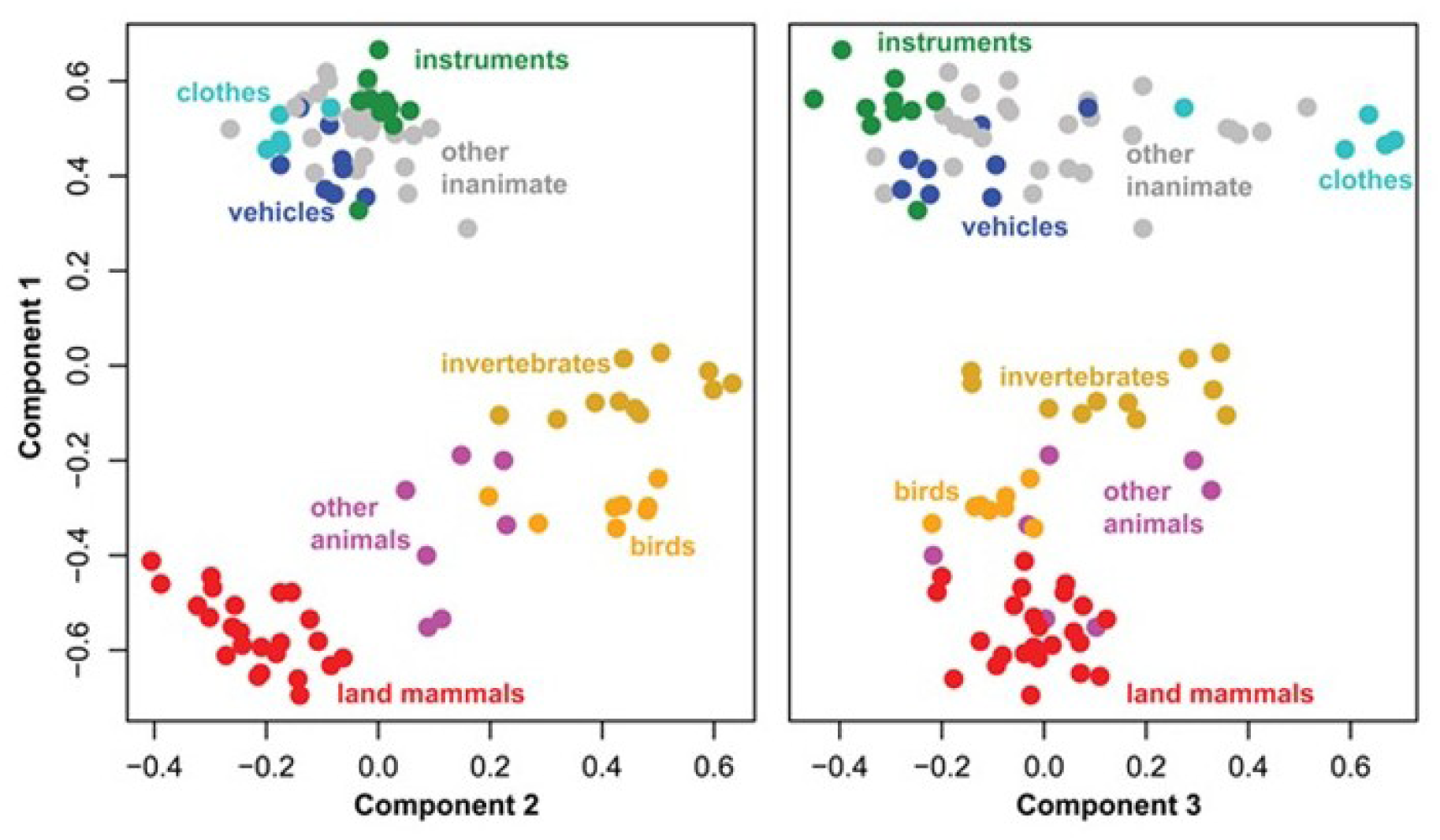
Coordinates of all stimuli on each target semantic dimension. Colours indicate category membership – land mammals (red), birds (orange), invertebrates (mustard), other animals (magenta), instruments (green), clothes (light blue), vehicles (dark blue), and other inanimate objects (grey). Reprinted with permission from Cox et al. ^16^.

The second dimension (4.4% of the variance) differentiates subclasses of animals, with mammals on one end and birds/invertebrates on the other; it is compatible with independent features that discriminate mammals from other animals (e.g., *has fur*, *can fly*, etc.) or a direction orthogonal to animacy within an inchoate semantic vector space. The third dimension (4.0% of the variance) largely differentiates manmade objects, with vehicles and instruments on one end, clothing on the other, and various miscellaneous items in between. It is compatible with independent features that differentiate kinds of artifacts (e.g., *is hard* vs. *is soft*) or as a third orthogonal direction within an inchoate semantic vector space.

RSL models were fitted using LASSO regularization (which assumes that the neural code is sparse and is indifferent to whether features encode one or multiple dimensions) or using grOWL regularization (which additionally assumes that important features encode multiple dimensions simultaneously).

In all cases models were fitted and evaluated using ten-fold nested cross-validation with an inner loop used to find optimal hyperparameters and with the resulting model then assessed on an outer-loop hold-out set of unseen items. For classification, performance was assessed as the mean classification accuracy over hold-out folds for classification models. For RSL, the Pearson correlation between predicted and true coordinates for held-out items was calculated for each hold-out fold and the mean of these correlations was used as the performance metric. To test for significance, the mean accuracy or correlation across participants was evaluated against an expected null distribution computed via bootstrapped permutations (see Methods; Stelzer et al., 2013).

### ATL analyses

Regions of interest are shown in Figure 2 and results for binary classification are shown in Figure 2A and Table 2. All models showed above-chance classification on held-out items, indicating that animacy information is encoded in both hemispheres, and in both anterior and posterior extents of the vTL — a pattern inconsistent with domain-general pointer representations. Classification with SOS-LASSO was reliably better than with LASSO for all models except right anterior vTL, suggesting that the regions encoding animacy information within vTL are anatomically clustered.

**Figure 2:**
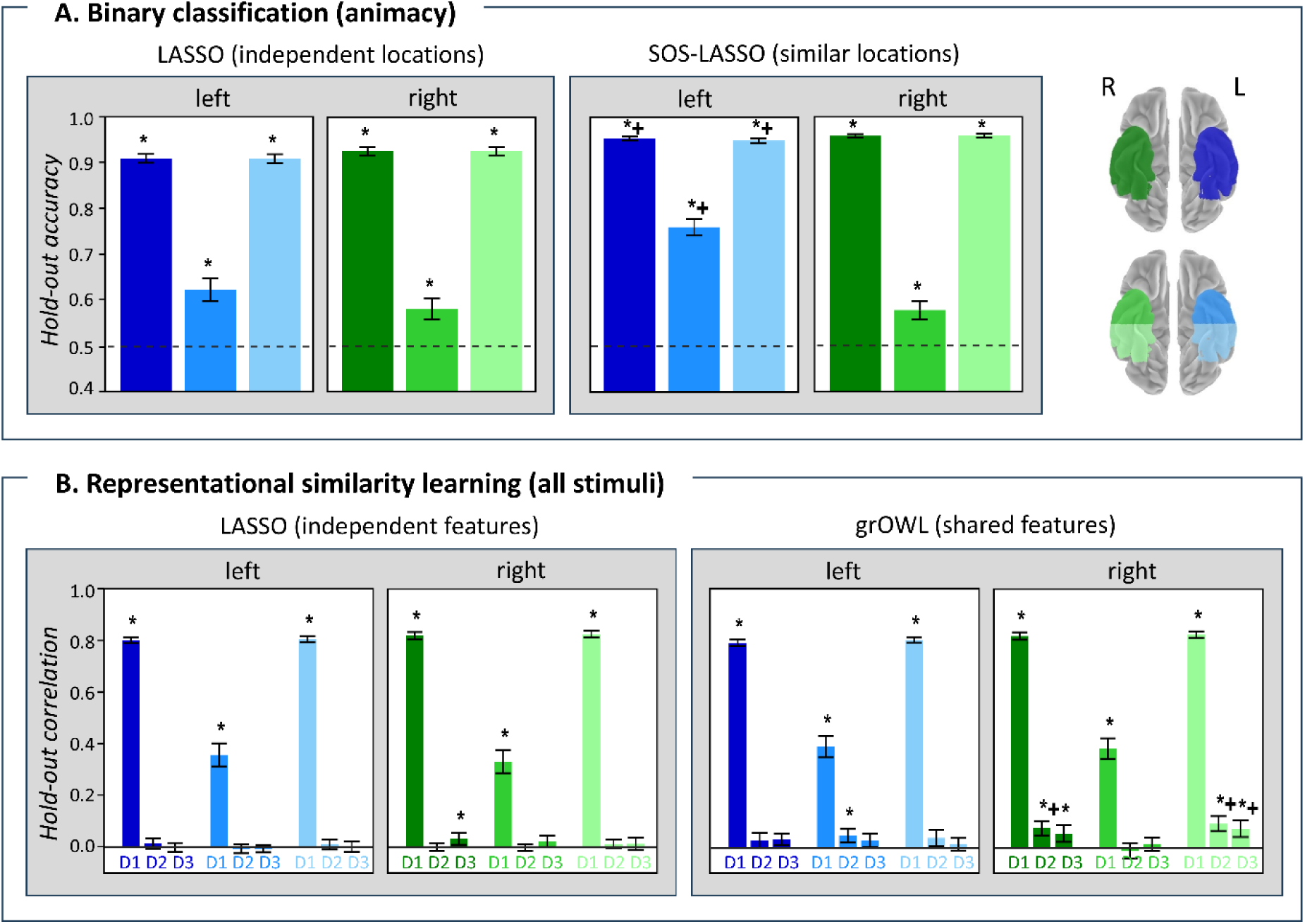
Decoding results in regions of interest. Surface plots (top right) indicate which bar corresponds to which ROI or half-ROI. (A) Mean and 95% confidence interval of the hold-out accuracy for regularised logistic regression classifiers trained on beta values to discriminate animate from inanimate items. Classifiers were trained using either LASSO or SOS-LASSO regularization separately for each ROI. Stars (*) indicate accuracy significantly better than expected under the null hypothesis (as determined by permutation test, probabilities adjusted to control the false-discovery rate at α = 0.05). Crosses (+) indicate a significant difference between classifier accuracy using that regularization penalty and accuracy in the same ROI using the other regularization penalty (probabilities adjusted to control the false-discovery rate at α = 0.05). (B) Mean and 95% confidence interval of the hold-out correlation for RSL models trained on beta values to predict the coordinates of held-out stimuli on three target semantic dimensions (dimension 1, D1; dimension 2, D2; and dimension 3, D3). Models were trained separately for each ROI using either LASSO or grOWL regularization fitted separately to each dimension or grOWL regularization fitted to all three dimensions simultaneously. Stars (*) indicate a significant correlation between predicted and target coordinates (as determined by permutation test, probabilities adjusted to control the false-discovery rate at α = 0.05). Crosses (+) indicate a significant difference between correlation on that dimension using that regularization penalty and correlation on the same dimension in the same ROI using the other regularization penalty (probabilities adjusted to control the false-discovery rate at α = 0.05).

**Table 2:**
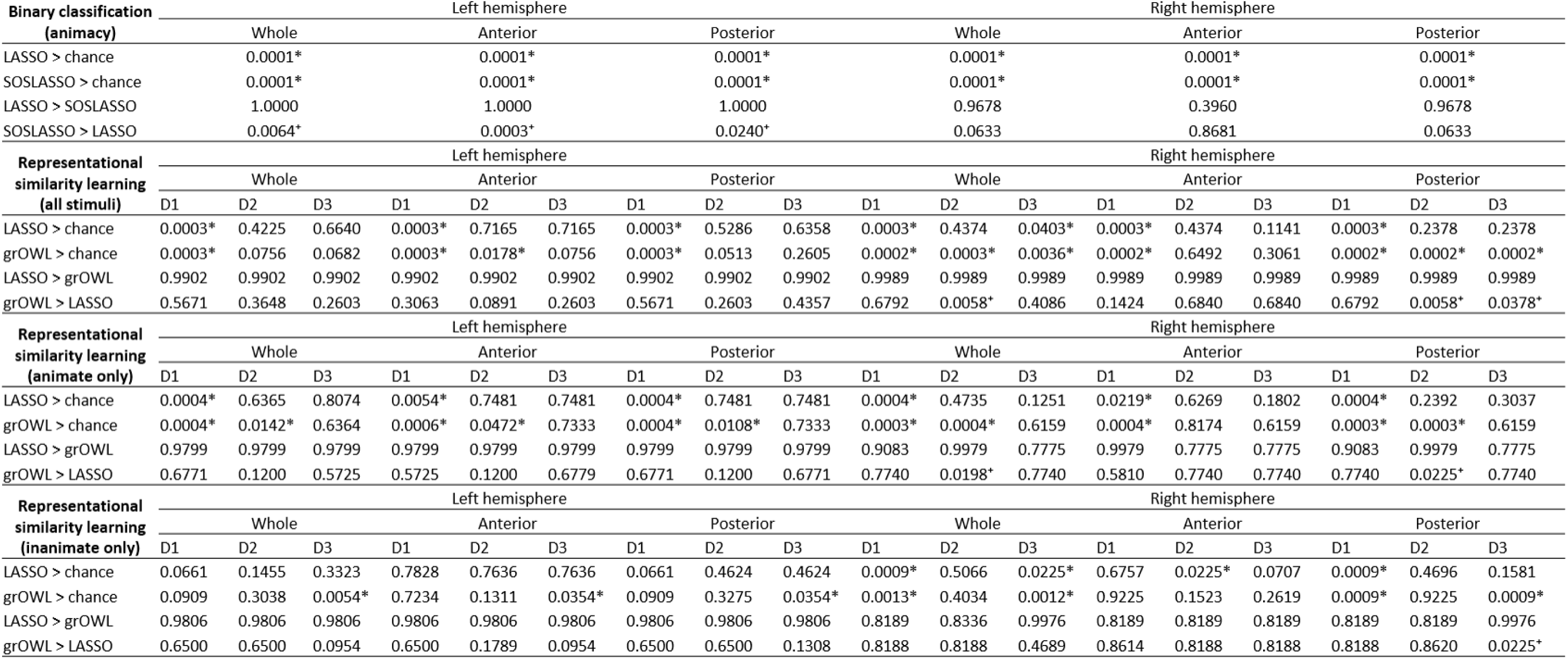
Permutation p-values for all hold-out correlations shown in Figure 1. Stars (*) indicate a significant correlation between predicted and target coordinates (as determined by permutation test, probabilities adjusted to control the false-discovery rate at α = 0.05). Crosses (+) indicate a significant difference between correlation on that dimension using that regularization penalty and correlation on the same dimension in the same ROI using the other regularization penalty (probabilities adjusted to control the false-discovery rate at α = 0.05).

For RSL, Figure 2B shows the mean hold-out correlation between predicted and true coordinates on each target dimension for each model, calculated across all stimuli. Table 2 shows the statistical results. All models showed very good decoding of the first target dimension in both hemispheres and all ROIs. Since the first dimension separates animate and inanimate items, this is consistent with the binary classification result. Models fitted with both regularization penalties produced significant correlations on at least one other dimension that, though small relative to the first component, were significantly higher than expected by chance and similar in magnitude to those observed in representational similarity analyses ^50^ (dimension 3 for LASSO, and both dimensions 2 and 3 for grOWL). When fitted using all vTL voxels, models regularised with grOWL showed reliable decoding of all three semantic dimensions from right-hemisphere activation, and in many cases the correlations were greater than those obtained with LASSO, which suggests that features encode multiple dimensions simultaneously.

Figure 3 shows predicted/true correlations calculated separately for animate and inanimate items in order to test for decoding of within-domain structure. Table 2 shows the statistical results. For animate items, models fitted with LASSO only performed above chance for dimension 1 in all ROIs. In contrast, models fitted with grOWL reliably decoded both the first and second dimensions in both hemispheres, in the posterior halves of each, and in the anterior half of the left hemisphere – a result that is incompatible with representations that are both pointer-based and animal-specific.

**Figure 3:**
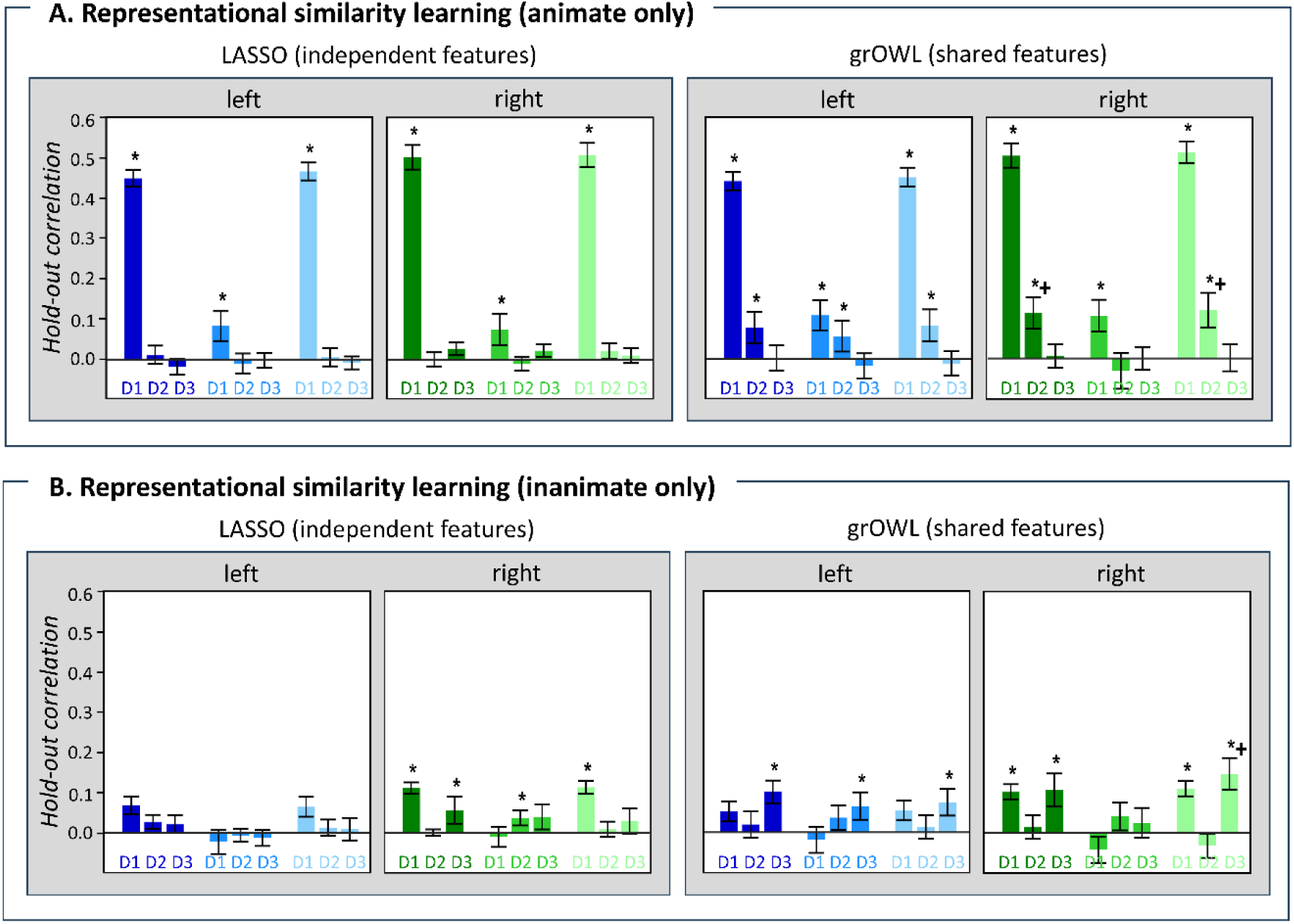
Hold-out correlations within-domain in regions of interest. Mean and 95% confidence interval of the hold-out correlation for RSL models trained on beta values to predict the coordinates of held-out stimuli on three target semantic dimensions (dimension 1, D1; dimension 2, D2; and dimension 3, D3). Models were trained using either LASSO or grOWL regularization separately for each ROI. Stars (*) indicate a significant correlation between predicted and target coordinates (as determined by permutation test, probabilities adjusted to control the false-discovery rate at α = 0.05). Crosses (+) indicate a significant difference between correlation on that dimension using that regularization penalty and correlation on the same dimension in the same ROI using the other regularization penalty (probabilities adjusted to control the false-discovery rate at α = 0.05). (A) Hold-out correlations for animate items only; (B) Hold-out correlations for inanimate items only.

For inanimate items, LASSO showed no reliable decoding in the left hemisphere and a somewhat mixed pattern in the right. In contrast, grOWL showed reliable decoding of dimension 3 in all three left-hemisphere ROIs, and of dimensions 1 and 3 in the right hemisphere for both the posterior half and the whole-vTL ROIs. This result indicates that representations within the vTL are not animal-specific. Again, the larger correlations obtained with grOWL suggest that features encode not one but multiple dimensions.

Note that, as shown in Figure 1, dimension 2 differentiates animals but not inanimate items, while the reverse is true for dimension 3. Accordingly, the decoding models generally showed reliable correlations within-domain on dimension 2 for animate items and on dimension 3 for inanimate items.

### Key observations

In summary, the reliable decoding of graded within-domain semantic structure for both animate and inanimate concepts is inconsistent with theories proposing that the ATL represents semantic information *only* within the animal domain. This graded result is also incompatible with all pointer-based theories. RSL models that preferred to predict all three dimensions from the same voxel subsets (grOWL regularization) often performed better than those with no such preference (LASSO), including the decoding of dimension 2 within animates and dimension 3 within inanimates from right-hemisphere activation. Notably, models regularised with LASSO never showed reliably better decoding than those regularised with grOWL. These findings are inconsistent with the view that separate neural populations encode independent semantic features, but consistent with the alternative view that the ATL encodes an inchoate semantic vector space in which each neural population contributes to the representation of multiple semantic dimensions. Finally, regularization with SOS-LASSO significantly improved binary classification in almost all ROIs, suggesting that the location of animacy information within the ATL is relatively contiguous within and consistently localised across individuals. Together, the only hypothesis consistent with the full pattern of results is that both anterior and posterior regions of the ATLs use anatomically clustered neural populations to encode an inchoate, domain-general semantic vector space (Hypothesis 9 in Table 1).

### Whole-cortex analyses

Figure 4A and Table 3 show the binary classification results. Accuracy was extremely high for classifiers fitted with LASSO and SOS-LASSO (hold-out accuracy ≅ 0.95). Accuracy did not differ reliably between these approaches, possibly due to ceiling effects.

**Figure 4:**
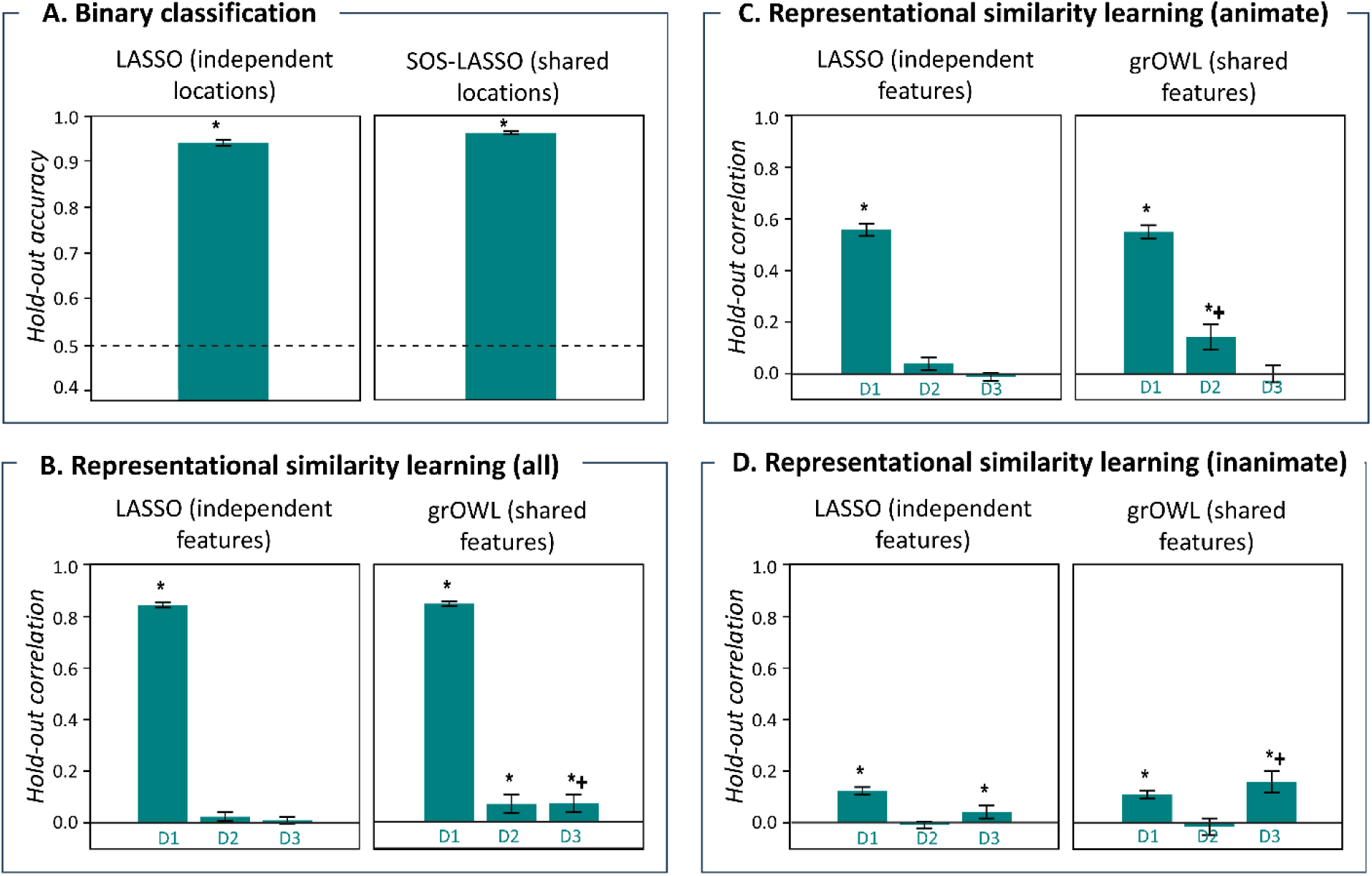
Decoding results at the whole-brain level. (A) Mean and 95% confidence interval of the hold-out accuracy for regularised logistic regression classifiers trained on beta values to discriminate animate from inanimate items. Classifiers are trained using either LASSO or SOS-LASSO regularization. Stars (*) indicate accuracy significantly better than expected under the null hypothesis (as determined by permutation test, probabilities adjusted to control the false-discovery rate at α = 0.05). (B) Mean and 95% confidence interval of the hold-out correlation for RSL models trained on beta values to predict the coordinates of held-out stimuli on three target semantic dimensions (dimension 1, D1; dimension 2, D2; and dimension 3, D3). Models are trained using either LASSO or grOWL regularization. Stars (*) indicate a significant correlation between predicted and target coordinates (as determined by permutation test, probabilities adjusted to control the false-discovery rate at α = 0.05). Crosses (+) indicate a significant difference between correlation on that dimension using that regularization penalty and correlation on the same dimension using the other regularization penalty (probabilities adjusted to control the false-discovery rate at α = 0.05). (C) Hold-out correlation for animate items only; (D) hold-out correlation for inanimate items only.

**Table 3:** Permutation p-values for all hold-out correlations shown in Figure 4. Stars (*) indicate a significant correlation between predicted and target coordinates (as determined by permutation test, probabilities adjusted to control the false-discovery rate at α = 0.05). Crosses (+) indicate a significant difference between correlation on that dimension using that regularization penalty and correlation on the same dimension in the same ROI using the other regularization penalty (probabilities adjusted to control the false-discovery rate at α = 0.05).

| **Binary classification**<br>**(animacy)** |  |  |
| --- | --- | --- |
| LASSO > chance |  | 0.0001* |
| SOSLASSO > chance |  | 0.0001* |
| LASSO > SOSLASSO |  | 0.8513 |
| SOSLASSO > LASSO |  | 0.1488 |

| **Representational**<br>**similarity learning**<br>**(all stimuli)** |  |  |  |
| --- | --- | --- | --- |
|  | D1 | D2 | D3 |
| LASSO > chance | 0.0003* | 0.1143 | 0.2682 |
| grOWL > chance | 0.0003* | 0.0004* | 0.0004* |
| LASSO > grOWL | 0.9951 | 0.9951 | 0.9951 |
| grOWL > LASSO | 0.4846 | 0.0636 | 0.0150* |

| **Representational**<br>**similarity learning**<br>**(animate only)** |  |  |  |
| --- | --- | --- | --- |
|  | D1 | D2 | D3 |
| LASSO > chance | 0.0003* | 0.0625 | 0.7556 |
| grOWL > chance | 0.0001* | 0.0001* | 0.4906 |
| LASSO > grOWL | 0.9146 | 0.9938 | 0.9146 |
| grOWL > LASSO | 0.5459 | 0.0189* | 0.5459 |

Table 3: Permutation p-values for all hold-out correlations shown in Figure 4.
| **Representational**<br>**similarity learning**<br>**(inanimate only)** |  |  |  |
| --- | --- | --- | --- |
|  | D1 | D2 | D3 |
| LASSO > chance | 0.0003* | 0.6160 | 0.0282* |
| grOWL > chance | 0.0010* | 0.6624 | 0.0003* |
| LASSO > grOWL | 0.6265 | 0.6265 | 0.9995 |
| grOWL > LASSO | 0.6347 | 0.6347 | 0.0018* |

Figure 4B and Table 3 show results for RSL. Models fitted with LASSO and grOWL showed very good decoding of the first target dimension (r ≅ 0.8), but only models fitted with grOWL showed reliable decoding of the second and third dimensions. Note that both LASSO and grOWL prefer sparse solutions, so neither is likely to have discovered the entire semantic code. Nevertheless, this result suggests that, across the cortex, the features that best predict multidimensional semantic structure do so via an inchoate vector-space code in which neural populations conjointly of multiple semantic dimensions.

Figures 4C and 4D and Table 3 show hold-out correlations calculated separately for animate vs. inanimate items. Both approaches achieved reliable decoding of within-domain structure on dimensions 1 and 2 for animate items, and on dimensions 1 and 3 for inanimate items. Within-domain decoding was, however, reliably better for grOWL than LASSO on dimension 2 for animates and dimension 3 for inanimates, again suggesting a conjoint, vector-space semantic code.

To visualise the anatomical locations of voxels contributing to decoding performance, we first used permutation testing to evaluate which voxels were selected by a decoding model more often than would be expected if no relationship exists between voxel activation and outcomes (see Methods). From these results we generated the cortical surface group maps shown in Figures 5 and 6. In these plots, coloured areas show cortical regions selected by a decoding method reliably more often than expected under the permutation null distribution. The colour indicates the proportion of decoding model coefficients that were positive across all participants, which illustrates how an increase in BOLD corresponds to semantic information encoding. Regions shown in blue show received mostly *positive* coefficients across participants; those in orange/red received mostly *negative* coefficients. Because animals were coded as 0 and inanimate objects as 1, for the classification analysis (Figure 5), increased activity in an orange/red region is associated with increased probability that the stimulus is animate and increased activity in a blue region is associated with increased probability that the stimulus is inanimate. For the RSL analysis (Figure 6), increased activity in an orange/red regions is associated with a lower coordinate and increased activity in a blue region is associated with a higher one (see Figure 1). In both analyses, regions in green were important for decoding but received mixed positive and negative coefficients across participants.

**Figure 5:**
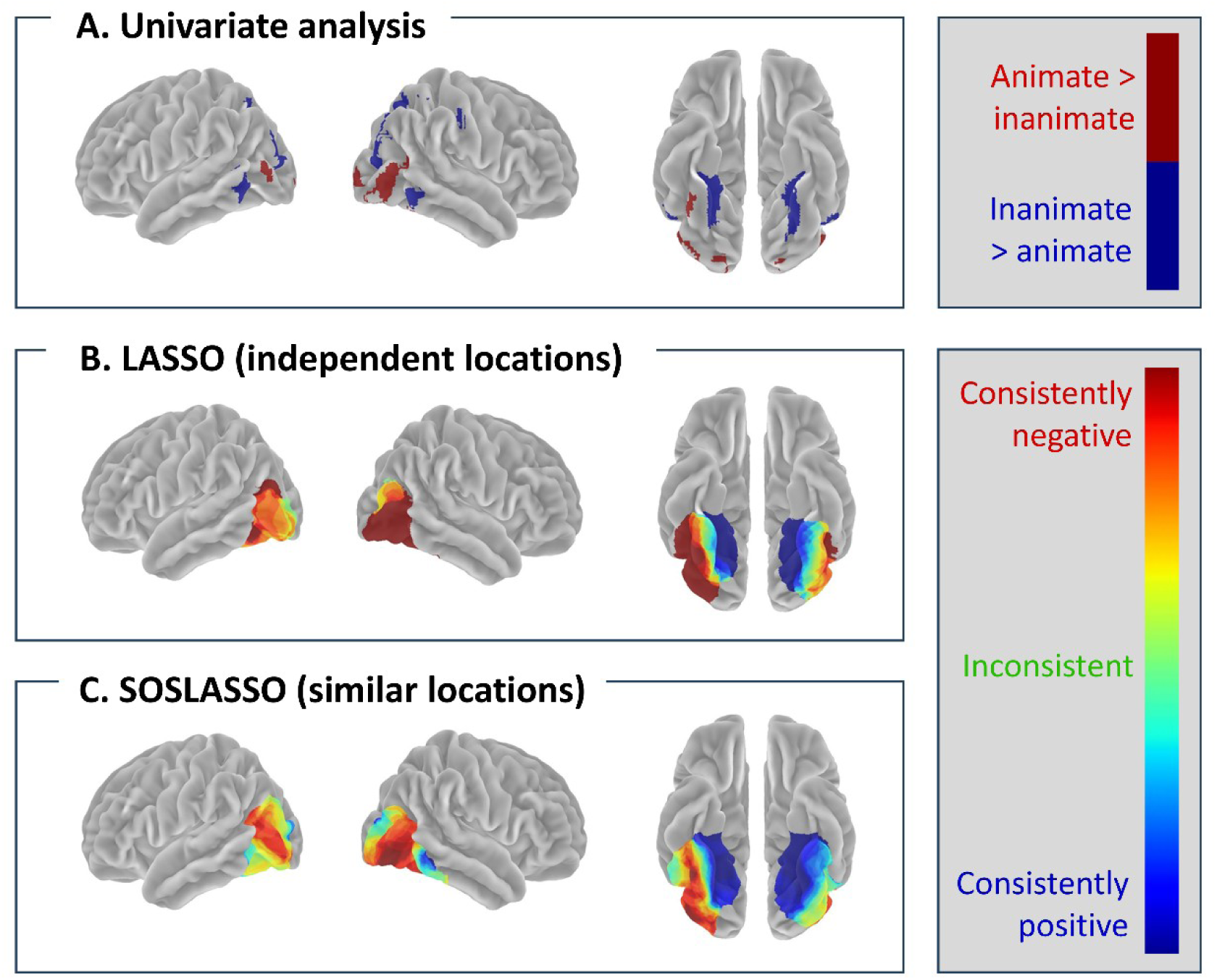
Coefficient maps for classification. Values are shown projected to the surface (fsaverage template). (A) Univariate effects (A>I, red; I>A, blue) are shown projected to the cortical surface and are cluster-corrected at p < 0.001. (B) Coefficients for logistic regression classifiers trained with LASSO regularization on beta values to discriminate animate from inanimate items. Coloured areas show voxels selected more often than expected from a null distribution computed via permutation testing and controlling the false discovery rate at α = 0.05 (see Methods). Colours indicate the proportion of coefficients that are positive across participants, with coloured labels indicating what an increase in activation signifies for a given decoder. For classification, cool colours indicate areas where elevated activation systematically signifies inanimate items across participants; warm colours indicate the reverse; and green indicate regions that contribute significantly to prediction, but in different directions across participants. (C) Coefficients for logistic regression classifiers trained with SOS-LASSO regularization.

**Figure 6:**
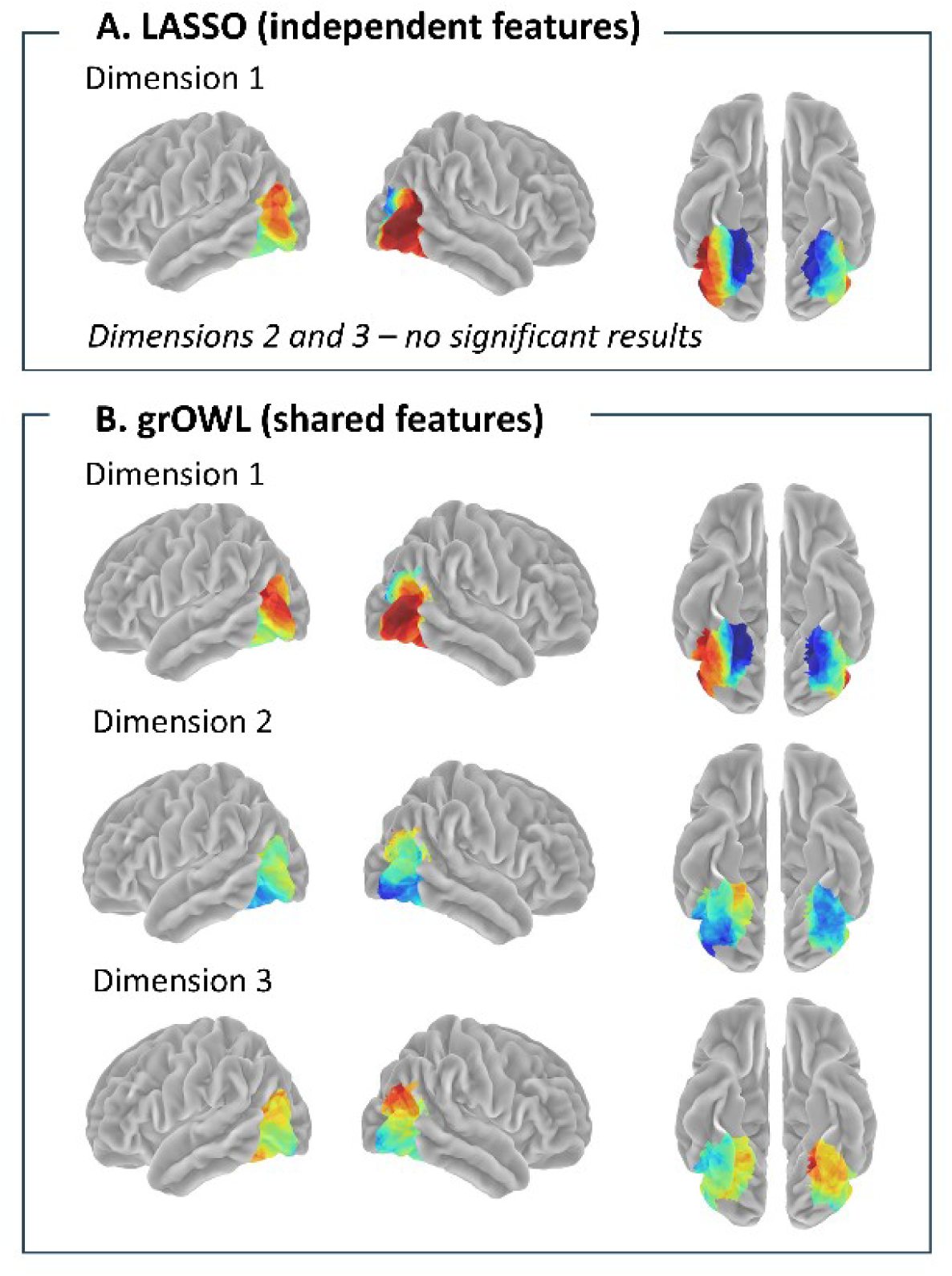
Coefficient maps for RSL. Values are shown projected to the surface (fsaverage template). (A) Coefficients for RSL models trained with LASSO regularization on beta values to predict the coordinates of held-out stimuli on three target semantic dimensions. Coloured areas show voxels selected more often than expected from a null distribution computed via permutation testing and controlling the false discovery rate at α = 0.05 (see Methods). Hues indicate the proportion of coefficients that are positive across participants, with coloured labels indicating what an increase in activation signifies for a given decoder. For dimension 1, cool colours indicate areas where elevated activation systematically signifies inanimate items across participants; warm colours indicate regions where elevated activation signifies animate items (for dimensions 2 and 3, no coefficients reached significance). (B) Coefficients for RSL models trained with grOWL regularization. Cool colours signify regions where elevated activation reliably signifies inanimate items (on dimension 1), birds or invertebrates (on dimension 2) or clothes (on dimension 3); warm colours indicate regions where elevated activation signifies animals (on dimension 1), mammals (on dimension 2) or instruments or vehicles (on dimension 3).

All models selected the same broad regions of ventral and lateral temporal and occipitotemporal cortex. Coefficients from binary classifiers (Figures 5B, 5C, 6A and 6B) and from the first dimension of the grOWL-based RSL model all showed a remarkably similar pattern – elevated activation in ventromedial temporal lobe bilaterally signified an inanimate stimulus, and elevated activation in more ventrolateral and lateral occipitotemporal regions signified an animate stimulus. Univariate results (Figure 5A) showed the same pattern but failed to detect regions which in which the coefficient direction differed across participants (green regions in panels B and C).

In Figure 6B, RSL coefficients for dimensions 2 and 3 occupied the same anatomical regions, consistent with grOWL’s preference for solutions in which the same voxels predict all target dimensions. The RSL coefficients for dimensions 2 and 3 were mixed in direction across much of the selected regions. Areas that did show directional consistency across participants were anatomically situated somewhat differently than those for dimension 1. For dimension 2 (separating animals), elevated activation consistently signified birds or invertebrates in left posterior ventrolateral temporal cortex but signified mammals in a more medial and anterior left ventral temporal region. For dimension 3 (separating artifacts), elevated activation in right ventromedial temporal cortex and right temporo-parietal junction consistently signified a greater likelihood of instruments or vehicles than clothing. While we do not wish to attribute too much significance to these exact patterns of coefficient distributions, the results do clearly indicate that (a) the same cortical regions encode, not just animacy, but also graded information about semantic similarity along multiple orthogonal semantic dimensions, both within and between domains, and (b) that the neuroanatomical consistency and organization of this information across the implicated regions differs for different semantic dimensions.

## 3. Discussion

Most multivariate studies of semantic representation employ a single kind of analysis, and so are constrained by that pipeline’s assumptions about the neural code and limited in the kinds of representational coding that they can and cannot detect ^11^. We considered a comparative multivariate decoding approach that, in exploiting the strengths and limitations of diverse methods, clarified *what*, *how*, and *where* semantic information is encoded in the ATL and elsewhere. Our results suggest that, given drawings of familiar objects as input, the ATL encodes semantic information across conceptual domains via an inchoate multidimensional vector-space with important neural populations anatomically clustered within and across participants (Hypothesis 9 in Table 1), and that the posterior temporal and occipitotemporal cortices encode similar information via a similar vector-space code.

Regarding the *domain* of representation (the “what” question), RSL successfully decoded multidimensional structure within and between both animate and inanimate items for both anterior and posterior ROIs, disconfirming the hypothesis that ATL representations are animal-specific. This finding aligns with results from neuropsychology, which demonstrate that patients with atrophy of the ATL do not show category-specific impairments ^36,51^; with results from neuro-computational modelling, which show a similar domain-general impairment following damage to a hub that intermediates between modalities ^2,4,52^; with results from neurostimulation, which show transient domain-general semantic impairment following ATL stimulation ^18^; and with univariate neuroimaging studies, which implicate the ATL in the representation of multiple domains ^26,53^. Note that a domain-general hub does not preclude the possibility of category-specific deficits - neuro-computational modelling also demonstrates that, if damage to a domain-general hub is followed by a period of recovery, a category-specific pattern of semantic impairments can appear, resembling those observed in patients with herpes simplex virus encephalitis (HSVE)^52^.

Regarding the *nature* of representation within the ATL (the “how” question), successful classification by animacy disconfirms domain-general pointer-based hypotheses—suggesting that, rather than mediating structured representations encoded elsewhere in the cortex, the ATL computes structured representations of its own ^33^. This result accords with prior simulation work suggesting that acquiring representations that express conceptual structure requires a system capable of re-representing the sensory-motor structure expressed within modality-specific regions to capture trans-modal similarities^4,38^. Whether this re-representation is composed of independent features, or of populations that conjointly encode an inchoate vector space, cannot be resolved simply by considering which semantic dimensions can be decoded—the successful decoding of semantic dimension 2, for instance, could arise under both feature-based and inchoate-vector-space hypotheses. Rather, it is the comparative decoding performance across different regularization techniques that provides the key information: because grOWL regularization yields better decoding than LASSO, this suggests that the ATL encodes semantic information via inchoate vector-space representations and not features (because grOWL encourages solutions that predict multiple target dimensions from the same neural features whereas LASSO fits each decoded dimension independently).

Finally, the comparison of different regularisers likewise illuminates the localisation of the ATL signal that encodes semantic information (the “where” question). Models constrained to encourage anatomical similarity (SOS-LASSO) out-perform those that ignore such information (LASSO), supporting ATL regions encoding semantic information somewhat anatomically contiguous within and consistently localised across individuals. To our knowledge, no prior work bears directly on this question. While the well-known hub-and-spoke model suggests the ATLs encode a domain-general, inchoate semantic vector space^1,2,36,38^, it has little to say about the anatomical distribution of signal-carrying neural populations within and across individual ATLs. In shedding light on this question, the current results offer new information that may help to refine neuro-computational models and to better understand patterns of impaired semantic behaviours following ATL injury.

The pattern of results for whole-cortex decoding was similar in many respects: classification by animacy was strongly reliable, and RSA showed successful decoding of within-and between-domain similarities across all three semantic dimensions, with grOWL regularization generally performing better than LASSO. The main difference was that we did not observe an advantage in animacy classification SOS-LASSO relative to LASSO—both methods showed very high decoding accuracy (∼0.95) from features selected in the mid to posterior extent of the ventral temporal lobes and in the lateral occipital regions bilaterally. The equivalent classification accuracy for the two methods may reflect a ceiling effect. A more sensitive test of within-and across-participant consistency in localization will require a comparison of regularisers that implement SOS-like anatomical constraints while capable of decoding more graded / less robust semantic information (like RSA).

All whole-brain analyses found semantic structure encoded in the posterior temporal and occipitotemporal cortices bilaterally. Prior work proposed that these regions may contain pointers for inanimate objects ^27,54,55^ or independent visual features ^56^; but successful decoding of multidimensional semantic structure is incompatible with the first of these hypotheses and the superior performance of grOWL relative to LASSO is incompatible with the second. Instead, this pattern again suggests a distributed semantic code in which voxels across the selected regions jointly express graded, multi-dimensional semantic structure amongst perceived images.

We note, however, that the surface rendering of model coefficients for decoding the first semantic dimension (animacy) shows clear anatomical structure: greater activation in medial posterior ventro-temporal regions bilaterally signals inanimate items, while greater activation in lateral posterior ventro-temporal regions and in lateral occipital cortex bilaterally signals animate items. Critically, however, the *same regions* also simultaneously express orthogonal semantic structure—dimensions 2 and 3—via a neuro-semantic code with a different anatomical organization (Figure 6B). Moreover, grOWL regularization yielded better decoding accuracy for these additional dimensions, suggesting that the additional semantic structure is encoded across the same voxels also expressing animacy information. Thus it may not be useful to view these regions as dedicated solely to distinct, local representations of different semantic domains, but as jointly encoding multiple other semantic dimensions simultaneously with animacy.

The whole-brain analysis did not select voxels within the ventral ATL; and yet the ROI-based analyses established that these regions do encode graded, multi-dimensional semantic structure. The disjunct makes it clear that the regularised regression approaches we have considered do not fully lay bare all neural systems expressing semantic information. These approaches all rely on some form of sparsity as a vehicle for voxel selection. Other common approaches such as ridge regression or elastic net spread small weights across all voxels and thus make it difficult to determine which voxels are important for the decoder performance (Rogers & Cox, XX) . In contrast, sparse regularisers pressure many coefficients to zero and thus draw a clear line between “selected” and “unselected” voxels; but even signal-carrying voxel sets can receive zero coefficients if the information they express is already accounted for by other selected voxel sets. While structured-sparse penalties like grOWL and SOS-LASSO compensate for this tendency to some degree (see XX), the failure of the whole-brain analysis to select ATL regions establishes that both approaches can still miss reliable signal. Thus, while cross-validated model performance establishes that selected regions do encode semantic structure, such structure may also be encoded within other neural systems not selected in the whole-brain analysis. Supplementary Figures S1-S3 describe additional analyses suggesting why ATL regions in particular may be dropped by sparse whole-brain regularization.

Finally, our analyses self-evidently characterise the neural code at the level of *voxels*. They do not address questions about structure at finer spatial or temporal scales. Functional activation of voxels provides a particularly useful abstraction of neural activity for cognitive neuroscience that enables convergence of neuroimaging with computational models of cognitive function and with clinical neuropsychology. As one example, units in computational models behave in ways that more closely resemble the behaviour of voxels than that of single neurons^57,58^; as a second example, description of the neural code at the voxel level in healthy individuals facilitates noninvasive imaging studies of patients that aim to characterise representational change in disease^59^. Nevertheless, it is possible that characterizations of representational structure at finer spatial and temporal scales will reveal neural codes with properties that differ from those arising at the voxel level. The same comparative multivariate decoding approach developed here can be equally applied to such data in future work, potentially illuminating how local patterns of neuronal firing give rise to the voxel-level encodings we have documented here.

## 4. Conclusion

Multiple forms of neuroscientific data converge on the conclusion that the ATLs support semantic cognition ^1^. This work demonstrates that, out of 12 hypotheses about the domain, nature and location of representation in the ATLs, one hypothesis best explains these data – that the ATLs represent domain-general semantic information (*what*) via a vector-space code (*how*) that is anatomically clustered within and across individuals (*where*). Posterior temporal and occipitotemporal regions utilise a similar domain-general, vector-space code. Our results not only adjudicate competing hypotheses about semantic representation but also illustrate the power of the comparative multivariate approach – using the pattern of results across multiple decoding models to differentiate theories. The conjoining of theory with decoding analysis approach ^11^ has the potential to reveal how the brain represents, not just semantic knowledge, but any kind of information.

## 5. Methods

### 5.1. Participants

32 participants were recruited to the study. Of these, five were unable to complete the study either for medical reasons or because of technical problems during data acquisition. This left 27 participants (age range 19-50, mean age 27.96 years, 17 female, 10 male) who were all right-handed native speakers of British English, with normal or corrected-to-normal vision, and with no neurological or sensory disorders. All participants gave written informed consent and the research was approved by a local National Health Service (NHS) ethics committee.

### 5.2. Stimuli and task

Stimuli were 100 line drawings used in previous ECoG work ^16,45,46,60^ – 50 animals and 50 inanimate items including vehicles, clothes, musical instruments and a range of other objects ^61^ (https://github.com/slfrisby/7TConvergent/tree/main/stimuli/). Animate and inanimate items were matched for age of acquisition, visual complexity, familiarity, word frequency and name agreement ^45,46,62^. Particular care was taken to ensure that visual structure, which is known to correlate with semantic structure ^63^, could not drive a positive decoding result. In previous work using these stimuli^46^, logistic regression classifiers identical to those used in the main analysis were trained to discriminate animate from inanimate stimuli using using vectors describing visual structure, obtained with Chamfer matching. These classifiers did not perform significantly differently from chance, confirming that visual structure would not be able to account for a subsequent positive decoding result.

PsychoPy 2022.2.5 ^64^ was used to display stimuli and record speech. Stimuli were rear-projected onto a screen in the MRI scanner and observers viewed stimuli through a mirror mounted to the head coil directly above the eyes.

Participants were instructed to name each picture out loud as quickly and accurately as possible while keeping head motion to a minimum. To ensure that participants understood the instructions, they practiced the task before entering the scanner while the experimenter provided feedback about their head motion. Practice continued until the experimenter was satisfied that head motion was minimised.

Overt speech was recorded for each trial and responses were scored for accuracy with the study’s aim of decoding semantic similarity structure in mind. If the participant did not respond within the 4 seconds during which the stimulus was on-screen, the trial was marked as incorrect. If the participant responded incorrectly but named an object that the rater judged to be visually and semantically similar to the stimulus and at a similar level of specificity (e.g. “wasp” for bee), the trial was marked as correct; if the object that they named was semantically dissimilar or was named at a less specific level (e.g. “insect” for bee), the trial was marked as incorrect. It was important that neural responses did not reflect a “blend” of the semantics of multiple stimuli so, if the participant gave two synonyms (e.g. “gun, pistol”) the trial was marked as correct; if they named two different objects (“kettle, iron”) the trial was marked as incorrect even if one of the names was correct. Trials containing fillers such as “um” and “err” in addition to a correct response were marked as correct because the participants were neurologically healthy and there was no reason to believe that these utterances reflected anything other than normal speech processes. The mean and standard deviation of the accuracy scores were calculated.

The study had a fast event-related design with 4 runs per participant, collected in a single session. Each run began 16 s after the start of the MR sequence and consisted of 100 trials (each stimulus appeared once per run). Each trial consisted of a stimulus presented for 4 s followed by a fixation cross presented for 4 s. We chose not to jitter since we were aiming to model the height of the haemoodynamic response function (HRF) rather than its full shape and since our naming task made it almost impossible for the participant to guess what response would be required by the next trial. 4 sequences of animate and inanimate stimuli were created and counterbalanced across participants. The order of trials was optimised for the contrast between animate and inanimate stimuli using optseq2 (https://surfer.nmr.mgh.harvard.edu/fswiki/optseq2). For each run and each participant, a random order of animate stimuli and a random order of inanimate stimuli were created; the stimuli were then interspersed according to the sequence generated with optseq2.

### 5.3. Image acquisition

All images were acquired on a whole-body 7T Siemens MAGNETOM Terra MRI scanner (Siemens Healthcare, Erlangen, Germany) with an 8Tx32Rx head coil (Nova Medical, USA).

An MP2RAGE anatomical scan was acquired with the following parameters: 224 sagittal slices (interleaved acquisition), FOV 240 x 225.12 x 240 mm, voxel size 0.75 mm isotropic, TR 4300 ms, inversion times 840 and 2370 ms, TE 1.99 ms, flip angles 5° and 6°, GRAPPA acceleration factor 3, and duration 8 minutes and 50 seconds.

Next, manual B_0_-shimming was performed over the volume to be used for echo-planar imaging (EPI). This was necessary because of a known bug in the scanner software which renders automatic shims inaccurate. A double-echo steady state (DESS) sequence was used to generate a field map and the radiographer adjusted shim parameters manually with the aim of reducing the water linewidth, defined as full width of the spectrum at half height, to below 40 Hz. However, for some participants the adjustments proved time-consuming and so adjustment time was capped at 30 minutes from the start of scanning after which the best shim parameters were adopted. The actual average water linewidth was 38.80 Hz (standard deviation = 5.93 Hz).

All four functional runs of EPI used the multi-echo multiband sequence designed by Frisby et al. ^65^ to improve ATL signal quality. The sequence had the following parameters: 48 axial slices (interleaved acquisition), FOV 210 x 210 x 210 mm to cover the whole brain in most participants (visual inspection ensured that the ATL was included in the FOV for all participants and the FOV was tilted up at the nose to avoid ghosting of the eyes into the ATL), voxel size 2.5 mm isotropic (no gap), TR = 1510 ms, 3 echoes, TE_1_ = 11.80 ms, TE_2_ = 27.05 ms, TE_3_ = 42.30 ms, flip angle 63°, multiband factor 2, GRAPPA acceleration factor 3, phase partial Fourier 7/8, and A-P phase encoding direction. Immediately *before* each run, 5 additional volumes were acquired with the phase encoding direction changed to P-A to facilitate distortion correction during preprocessing.

### 5.4. Data analysis

All analysis code is available at: https://github.com/slfrisby/7TConvergent/.

#### 5.4.1. Preprocessing

For ease of data sharing we converted all DICOMs to BIDS format ^66^ using *heudiconv* (v1.1.1; ^67^).

Standard reproducible preprocessing pipelines designed for 3T-fMRI, such as *fMRIprep* ^68–70^ perform poorly on 7T-fMRI EPI data. Therefore, the analysis pipeline was split into stages using different software packages.

Since it is notoriously difficult to perform a good-quality brain extraction on MP2RAGE data (because of salt-and-pepper noise in the background and cavities), two pipelines were used. The MP2RAGE T1w (combined) image first had its background noise removed using O’Brien regularization ^71^ and was then submitted to CAT12 for segmentation ^72^; in SPM12 (https://www.fil.ion.ucl.ac.uk/spm/). The bias-corrected and global-intensity-corrected T1w (combined) image produced was provided to as input to the anatomy pipeline in *fMRIPrep* 21.0.1.The T1w (combined) image was corrected for intensity non-uniformity with *N4BiasFieldCorrection* ^73^ and skull-stripped with a Nipype implementation of the *antsBrainExtraction.sh* workflow (ANTs 2.3.3, https://github.com/ANTsX/ANTs; ^74,75^. Brain tissue segmentation of cerebrospinal fluid (CSF), white matter (WM) and grey matter (GM) was performed on the brain-extracted T1w (combined) image using *fast* (FSL 6.0.5.1, ^76^). Volume-based spatial normalization to standard space (MNI152NLin2009cAsym) was performed through nonlinear registration with *antsRegistration.sh* (ANTs).

Functional preprocessing was performed using in-house code, composed of functions from AFNI (v.18.3.03; ^77,78^; https://afni.nimh.nih.gov/), FSL (v.5.0.10; ^79–82^; https://fsl.fmrib.ox.ac.uk/fsl/fslwiki), tedana (v.24.0.0; ^83–86^; https://tedana.readthedocs.io/en/stable/index.html) and ANTS (v. 2.2.0; ^74,75^; https://github.com/ANTsX/ANTs). Slice timing was corrected to the middle slice using *3dTshift* (AFNI), motion was corrected with *3dvolreg* and *3dAllineate* (AFNI), using the first volume of the first run acquired as a reference (for each volume, the TE_1_ image was aligned and the resulting transform was applied to the TE_2_ and TE_3_ images), and skull-stripped using *BET* (FSL ^81^) to create a participant-specific brain mask. Then *tedana* was used to combine data from all echoes optimally based on T2* weighting ^87^ and to denoise the data using ICA. *tedana* conducts denoising by decomposing data using PCA and ICA, classifying components according to whether the signal scales linearly with TE (as BOLD signal does), and reconstruct the data using only BOLD-like components. The brain mask created with *BET* was used as the mask for this stage.

Next, unwarping was conducted using *topup* and *applytopup* (FSL; ^88^). Field displacement maps were calculated using ten volumes (five with A-P phase encoding direction, extracted from the start of each functional run, and five with P-A phase encoding direction, collected separately before each run) and the resulting correction was applied to all images. Finally, the mean EPI for each run was coregistered to the skull-stripped native structural image using a rigid-body registration in *AntsRegistrationSyN.sh* (ANTS). This produced transforms from native EPI space to native T1 space. For all decoding analyses, images remained in native space and were not smoothed. However, the transforms from native EPI to native T1 (for the first run) and the transforms from native T1 to standard space (MNI152NLin2009cAsym), with *antsApplyTransforms* (ANTS), were used to calculate the MNI-space coordinates that each voxel *would* have *were* images to be transformed into MNI space. These coordinates were used for visualization and were provided to classifiers trained with SOS-LASSO regularization, which take anatomical information as input. The transform from native EPI to native T1 space was also used to transform the participant’s grey-matter mask into native EPI space. For the univariate analysis, images were transformed into standard space (MNI152NLin2009cAsym) with *antsApplyTransforms* (ANTS) and smoothed with a 6 mm FWHM Gaussian filter in SPM12 (https://www.fil.ion.ucl.ac.uk/spm/).

#### 5.4.2. 1^st^-level (within-participant) GLM

Data were processed using the general linear model (GLM) approach implemented in SPM12 in MATLAB r2023b. For decoding, each stimulus was modelled as an individual boxcar function, 1.5 seconds in length because the stimulus in our stimulus set with the longest mean naming time had a mean naming time of 1323 ms ^61^ (periods between stimuli were modelled implicitly). These boxcar functions were subsequently convolved with SPM’s difference of gammas haemodynamic response function. The six motion parameters extracted during preprocessing were used as regressors of no interest. A single model was fitted for all runs, so an additional constant was included for each run. The micro-time resolution was set as the number of slices (n = 48), the micro-time onset was set as the reference slice for slice-time correction (n = 24), and the high-pass filter cutoff was 128 seconds. Each participant’s grey matter mask in native EPI space was used as a mask for the analysis. The parameter estimation method was restricted maximum likelihood estimation (ReML) and serial correlations were accounted for using an autoregressive AR(1) model during estimation. This produced one beta image per stimulus in native space. For univariate analysis, the same model was run on the normalised, smoothed data with identical parameters except that a grey matter mask in standard space (MNI152NLin2009cAsym) was used, that only two task regressors were used (animate stimuli named correctly and inanimate stimuli named correctly), and that animate stimuli named incorrectly and inanimate stimuli named incorrectly were modelled as additional regressors of no interest. The univariate contrasts of interest were greater activation for animate stimuli named correctly than inanimate stimuli named correctly (A > I) and greater activation for inanimate stimuli named correctly than animate stimuli named correctly (I > A). This produced contrast images in MNI space.

#### 5.4.3. Univariate analysis – 2^nd^-level (across-participant) GLM

Group-level t-tests of each contrast of interest (A > I, I > A) were conducted in SPM12. The group t-maps were assessed for significance by using a voxel-height threshold of p < 0.001 to define clusters and then a cluster-defining family-wise-error-corrected threshold of p < 0.05 for statistical inference. Group-level results were then projected to the pial surface (fsaverage template ^89,90^) using *-volume-to-surface-mapping* with trilinear interpolation in Connectome Workbench 1.5.0 and the thresholded maps were plotted on the pial surface using *plot_surf_stat_map* in *nilearn* implemented in Python 3.9.

#### 5.4.4. Multivariate decoding analysis

##### 5.4.4.1. Input data

To prepare data for decoding, beta images from all 4 presentations of each stimulus were averaged across runs. If a participant had named a stimulus incorrectly on one to three occasions, the beta image(s) corresponding to the incorrect trial(s) were not included in the averaging. If the participant named the stimulus incorrectly on all four trials, the values for each voxel were interpolated with the median value for that voxel over all other stimuli. This was done to ensure that each participant’s data would contain 100 items, maintaining a fold balance of five living and five nonliving stimuli and preventing decoding models from simply predicting the majority class.

For the vTL analysis, an ROI was constructed based on electrode coverage in a previous study ^45^; all electrode coordinates are available at https://github.com/slfrisby/7TConvergent/). Electrodes were converted to one NIFTI volume at native EPI resolution (2.5 mm isotropic) in MNI space and smoothed at 10 mm FWHM using SPM12. In order to exclude regions that lay outside the temporal lobe, an “exclusion mask” was created by taking the Harvard-Oxford cortical and subcortical atlases, removing the temporal lobe from the atlas, and adding the cerebellum from the AAL 2 atlas ^91^; since the Harvard-Oxford atlas does not include a cerebellum). Only voxels that lay *outside* this exclusion mask were included in the ROI. After masking the ROI was smoothed again at 4mm FWHM because, on inspection, this created a smooth shape that fully encompassed the temporal lobes. Since electrode coverage was far more extensive in the left than the right ROI, a mirror-image of the left ROI was used as the right ROI. The final ROIs in standard space (MNI152NLin2009cAsym) are shows projected to the surface in Figure 2 and are available at https://github.com/slfrisby/7TConvergent/tree/main/ROI/. Because the ROI was relatively large and we wished to achieve greater anatomical specificity, we also subdivided the ROI into anterior and posterior sections. Attempting to subdivide the ROI into distinct anatomical regions is challenging – the ATL was noted by Brodmann to feature gradual, rather than sharp, cytoarchitectural transitions^47^ – so we simply bisected it along the x-axis and analyzed the halves independently as well as the whole. The ROIs were backprojected into each participant’s native space using the transforms from native EPI to native T1 and the transforms from native T1 to standard space generate with *antsRegistration.sh*. Only beta values from voxels that overlapped with both the ROI and the participant’s native-space grey matter mask were provided as input to the decoding models.

For the whole-cortex analysis, only voxels that overlapped with the participant’s native-space grey matter mask were provided as input to the decoding models.

##### 5.4.4.2. Decoding models

We tested for semantic structure using two decoding methods. First, we used regularised logistic regression classifiers to decode binary animacy ^46^. One set of classifiers was trained using LASSO regularization ^92^, which assumes that the neural code is sparse (only a few of the features under consideration encode the target structure) and uncorrelated (important features exhibit uncorrelated activity). These classifiers were fitted using *glmnet* in MATLAB r2023b ^93^, using 3 – 0.2 as the range of possible lambda values. Each classifier took, as input features, beta values from multiple voxels from a single participant. A second set of classifiers was trained using SOS-LASSO regularization, which assumes that the neural code is relatively sparse but also in roughly the same anatomical location within and across participants. Full mathematical details, illustrating how SOS-LASSO incorporates these assumptions, can be found in Rao et al. ^94,95^ and Cox and Rogers ^41^. These classifiers were fitted using the WISC MVPA toolbox in MATLAB r2018b (https://github.com/crcox/WISC_MVPA/) with hyperband budget set to 25 (all other parameters were set as defaults, including a set size of 18 mm with 9 mm overlap and 3 – 0.2 as the range of possible lambda values). These classifiers took beta values from multiple voxels from multiple participants as input. Those data were used to find the best values of the hyperparameters lambda and alpha; then those hyperparameters were used to train a separate model, and find a separate set of coefficients, for each participant ^41,94,95^. Additionally, though data were in native space, the MNI-space coordinates that each voxel *would* have *were* images to be transformed into MNI space were also provided as input to the classifier.

Second, we used RSL to assess whether and where graded, multidimensional semantic structure was represented. Target semantic dimensions were modelled following the approach of Cox et al. ^16^ – details are provided in full there and are summarised here. Our starting point was a set of feature verification norms collected in a previous study ^48^ – participants freely listed semantic features of each item and the entire set of features, produced for *any* item, was then verified for *every* item (for example, a participant might list “breathes air” for a whale but not for a dog; in the final dataset, both a whale and a dog would be encoded as breathing air). First, a representational similarity matrix (RSM) was constructed that expressed the similarity between vectors of semantic features for each possible pair of stimuli. Next, singular value decomposition (SVD) was applied to extract three components accounting for 89.5% of the variance in the whole RSM (81.1%, 4.4%, and 4.0%, respectively). The coordinates of all 100 stimuli on each dimension are shown in Figure 1.

RSL models were fitted to predict the coordinates of each stimulus on the three target semantic dimensions from beta values from multiple voxels. One set of models used LASSO regularization and were trained using *glmnet* in MATLAB r2023b, using 6 – 0 as the range of possible lambda values. A second set of models used grOWL regularization, which assumes that the neural code is sparse but correlated – in other words, information is encoded by small groups of features that are correlated in their activity. Importantly, grOWL also assumes that the neural code is interdependent – features carry information about multiple dimensions simultaneously. Full mathematical details, explaining how grOWL incorporates these assumptions, can be found in Oswal et al. ^40^ and Cox et al. ^16^. grOWL models were trained using the WISC MVPA toolbox in MATLAB r2018a with the hyperband budget set to 25 (all other parameters were set as defaults, including 6 – 0 as the range of possible lambda values). All RSL models took, as input features, beta values from multiple voxels from a single participant.

##### 5.4.4.3. Cross-validation and model evaluation

For both logistic regression classifiers and RSL models, model performance was assessed via ten-fold nested cross-validation. Data were split into ten folds, each containing ten stimuli (five animate, five inanimate). In the first instance, stimuli were assigned randomly to the ten folds, but this order was then fixed so that models fitted with every kind of regularization were assessed using the same fold configuration. One fold was designated as the outer-loop hold-out set and a second fold was designated as the inner-loop holdout set. The stimuli in the remaining eight folds were used as training data to search for the best-performing hyperparameter(s). Models with these hyperparameter values were assessed on the inner-loop hold-out set. The folds were then reassigned so that ten stimuli previously used for training became the inner-loop hold-out set. Once all combinations of inner-loop hold-out set and training data had been explored, the best-performing hyperparameter value(s) were used to train a final model that was assessed on the outer-loop hold-out set. The procedure was then repeated with the nine other possible outer-loop hold-out sets.

The mean classification accuracy (for classifiers) or correlation between true and predicted coordinates (for RSL models) was calculated over all hold-out folds. Statistical significance was assessed via permutation testing. This was done for two reasons. First, since we used a fast event-related design rather than a slow event-related design, the haemodynamic response overlapped between trials. This means that, if temporally adjacent stimuli were divided between training and test sets, the stimulus in the training set may contain information about the stimulus in the test set and so may produce classification hold-out accuracies above 0.5 or hold-out correlations above 0. Second, cross-validated correlation (as was calculated for RSL models) is known to produce predicted coordinates that sometimes correlate *negatively* with the target coordinates in a hold-out set – the mean correlation under the null hypothesis may be below zero ^16,96^; therefore, the magnitude of the real correlation can be understood only with reference to the permutation distribution. In permutation testing, the stimulus labels were permuted before model training (the permutation orders were generated at random but then fixed so that the permutations for each model were comparable) and the procedure was repeated 100 times to produce 100 simulated accuracy or correlation values for each participant. A group-level empirical null distribution was simulated by randomly sampling one of the 100 values from each participant and computing a group average 10,000 times ^49^. One-tailed *p*-values were calculated (where *m* is the number of values in the permutation distribution and *b* is the number of values in the permutation distribution larger than the true value ^16,97^):

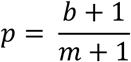

To test whether one model or ROI produced a higher accuracy or correlation than another, a group-level permutation distribution of differences was created using the differences between the values in both permutation distributions and using the group-level simulation procedure described above. A p-value was then calculated for the true difference. Statistical significance for all tests was defined after adjusting p-values to control the false discovery rate at α = 0.05 ^98^.

##### 5.4.4.4. Coefficient visualization

Ten-fold nested cross-validation does not provide a clear picture of which voxels are informative – each of the ten models selects its own set of coefficients ^41^. Therefore, to visualise coefficients, one model was trained for each participant using an inner loop of ten stimuli to identify the best hyperparameter(s) but no outer loop (i.e. all 90 stimuli that were not in the current inner loop hold-out set were used for hyperparameter exploration; therefore, all 100 stimuli were used for training). 100 permutation models were also trained for each participant (but no random sampling was conducted at the visualization stage). Coefficients were converted to NIFTI volumes at native EPI resolution (2.5 mm isotropic) in MNI space using SPM12, separately for each participant. Coefficients both for models trained on real data and for models trained on permuted data were projected to the pial surface (fsaverage template; ^89,90^) using *-volume-to-surface-mapping* with trilinear interpolation and smoothed on the surface at 6 mm FWHM using *–metric-smoothing* in Connectome Workbench 1.5.0. Two metrics were calculated – (1) the proportion of participants in which each vertex was selected and (2) the proportion of negative coefficients that each vertex received. (For classification, since animals were coded as 0 and inanimate objects as 1, a negative coefficient indicates that a positive beta value is associated with increased probability that the stimulus is animate. For RSL, although the dimensions capture more than just animacy, the same approach was taken for consistency.) The map of the proportion of negative coefficients was thresholded using the map of coefficient selection - vertices were included in the map only if they were selected significantly more frequently in the real distribution than they were in the permutation distribution, as assessed via binomial tests with p-values adjusted to control the false discovery rate at α = 0.05 ^98^. Finally, the thresholded map was plotted on the pial surface using *plot_surf_stat_map* in *nilearn* implemented in Python 3.9.

## Data and code availability

Data are available for scientific research purposes via the MRC Cognition and Brain Sciences Unit Data Repository (https://www.mrc-cbu.cam.ac.uk/publications/opendata/request/9274/). Code is publicly available at: https://github.com/slfrisby/7TConvergent/.

## Author contributions

Saskia L. Frisby – Conceptualization, Methodology, Investigation, Data Curation, Formal Analysis, Visualization, Writing – Original Draft, Writing – Review & Editing; Christopher R. Cox – Conceptualization, Methodology, Software, Writing – Review & Editing; Ajay D. Halai – Conceptualization, Methodology, Writing – Review & Editing, Supervision; Matthew A. Lambon Ralph – Conceptualization, Methodology, Writing – Review & Editing, Supervision; Timothy T. Rogers – Conceptualization, Methodology, Writing – Review & Editing, Supervision.

## Acknowledgements

We would like to thank the participants and the Wolfson Brain Imaging Centre radiographers.

