## Supplementary Materials 1: Additional Figures for "Comparative multivariate decoding adjudicates theories of semantic representation in the anterior temporal lobes and the rest of the cortex"

**Supplementary Materials S1: Additional Figures**

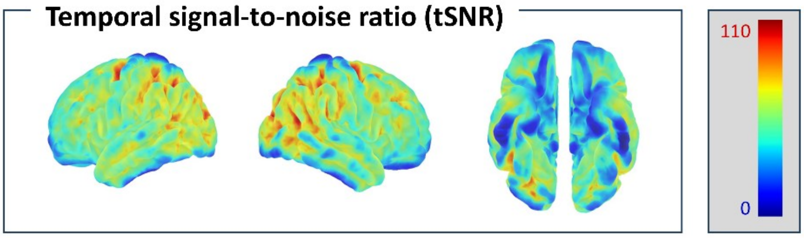


Supplementary Figure S1: Mean temporal signal-to-noise ratio. Values are shown projected to the surface (fsaverage template).

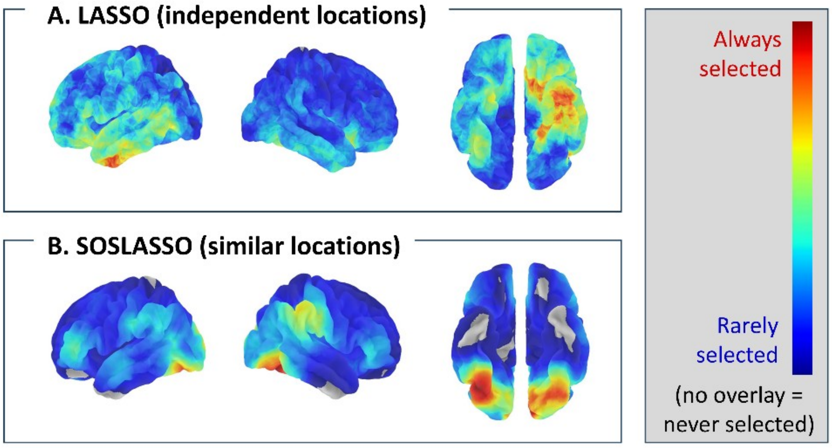


Supplementary Figure S2: Selection in the permutation distribution for classification. (A) Proportion of participants in which each vertex is assigned a non-zero coefficient by logistic regression classifiers trained with LASSO regularisation on *permuted* beta values to discriminate animate from inanimate items. Warm colours indicate that voxels in the region are selected more frequently. (B) Proportion of participants in which each vertex is assigned a non-zero coefficient by logistic regression classifiers trained with SOS-LASSO regularisation.

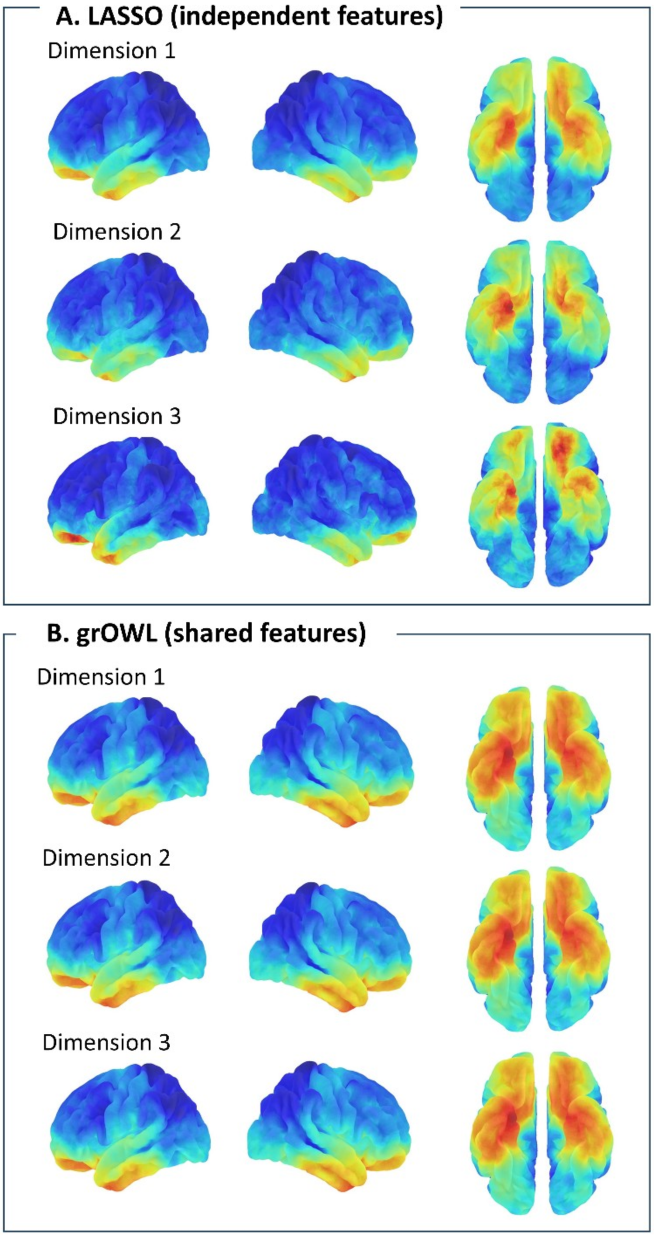


Supplementary Figure S3: Selection in the permutation distribution for RSL.(A) Proportion of participants in which each vertex is assigned a non-zero coefficient by RSL models trained with LASSO regularisation on *permuted* beta values to predict the coordinates of held-out stimuli on three target semantic dimensions. Warm colours indicate that voxels in the region are selected more frequently. (B) Proportion of participants in which each vertex is assigned a non-zero coefficient by RSL models trained with grOWL regularisation.
